# Evidence for widespread alterations in cortical microstructure after 32 hours of sleep deprivation

**DOI:** 10.1101/2021.06.22.449439

**Authors:** Irene Voldsbekk, Atle Bjørnerud, Inge Groote, Nathalia Zak, Daniel Roelfs, Ivan I. Maximov, Oliver Geier, Paulina Due-Tønnessen, Erlend Bøen, Yvonne S. Kuiper, Lise-Linn Løkken, Marie Strømstad, Taran Y. Blakstvedt, Bjørn Bjorvatn, Ulrik F. Malt, Lars T. Westlye, Torbjørn Elvsåshagen, Håkon Grydeland

## Abstract

Cortical microstructure is influenced by circadian rhythm and sleep deprivation, yet the precise underpinnings of these effects remain unclear. The ratio between T_1_-weighted and T_2_-weighted magnetic resonance images (T_1_w/T_2_w ratio) has been linked to myelin levels and dendrite density and may offer novel insight into the intracortical microstructure of the sleep deprived brain. Here, we examined intracortical T_1_w/T_2_w ratio in 41 healthy young adults (26 women) before and after 32 hours of either sleep deprivation (*n* = 18) or a normal sleep-wake cycle (*n* = 23). Linear models revealed significant group differences in T_1_w/T_2_w ratio change after 32 hours in four clusters, including bilateral effects in the insular, cingulate, and superior temporal cortices, comprising regions involved in attentional, auditory and pain processing. Across clusters, the sleep deprived group showed an increased T_1_w/T_2_w ratio, while the normal sleep-wake group exhibited a reduced ratio. These changes were not explained by in-scanner head movement, and 95% of the effects across clusters remained significant after adjusting for cortical thickness and hydration. Compared with a normal sleep-wake cycle, 32 hours of sleep deprivation yields intracortical T_1_w/T_2_w ratio increases. While the intracortical changes detected by this study could reflect alterations in myelin or dendritic density, or both, histological analyses are needed to clarify the precise underlying cortical processes.

## 1 Introduction

Insufficient sleep causes deficits in cognitive and affective processing[1], and is frequently reported among patients with neurodegenerative and psychotic disorders[2]. Chronic sleep deprivation is also common in mood and anxiety disorders [3]. Paradoxically, acute sleep deprivation has an intriguing antidepressant effect in some patients, peaking after around 32 hours[4, 5]. Findings from rodent studies suggest that acute sleep deprivation is linked to structural alterations in the cerebral cortex involving myelin and dendritic spines[6, 7]. However, we lack a clear understanding of the cortical mechanisms underlying sleep deprivation effects in humans.

Using magnetic resonance imaging (MRI) derived indices, several human studies have reported cortical changes following acute sleep deprivation[8, 9]. One night without sleep was linked to volume decreases in the insula and parietal cortex[8], and decreased cortical thickness in the medial parietal cortex[9]. However, the interpretation of the underlying mechanisms is limited by the macrostructural nature of these measures. Using microstructural imaging such as diffusion-weighted imaging (DWI), studies have reported sleep deprivation effects on white matter pathways[10, 11]. This includes alterations in radial diffusivity (RD), which has been related to myelin integrity[12]. DWI has also been applied to measure sleep deprivation effects in the cortex, showing declines in cortical mean diffusivity (MD)[13]. However, the neurobiological correlates of MD changes in the cerebral cortex are complex and not well understood.

Another index of intracortical microstructure is the ratio between T_1_-weighted and T_2_-weighted images (T_1_w/T_2_w ratio)[14]. Regional variation in T_1_w/T_2_w ratio has been found to relate to variation in histologically derived myelin levels[14, 15], and, recently, also to dendrite density[16]. While awaiting further histological comparisons, the T_1_w/T_2_w ratio might be sensitive to the very intracortical properties reported in rodents following sleep deprivation. Interestingly, a recent cross-sectional study reported associations between T_1_w/T_2_w ratio and self-reported sleep quality and sleep duration in several cortical regions[17]. While correlational, these findings may suggest that the T_1_w/T_2_w ratio is sensitive to sleep-related processes. Hence, testing whether acute sleep deprivation leads to alterations in the T_1_w/T_2_w ratio might help illuminate the underlying substrates of sleep deprivation effects in humans.

In this study, we investigated the effects of 32 hours of wake and sleep on cortical microstructure using the T_1_w/T_2_w ratio. In total, 41 healthy young adults underwent MRI before and after either sleep deprivation (*n* = 18) or a normal sleep-wake cycle (NSW, *n* = 23). As recent studies report time-of-day effects on other MRI modalities[9, 10, 13, 18-24], we aimed to control for such effects by comparing the effect of 32 hours of sleep deprivation with NSW. To this end, we first estimated within-subjects changes between the two MRI scans and then assessed whether these were significantly different between the sleep deprivation group and the NSW group. Based on (i) the links between sleep deprivation and myelin and dendritic spine alterations and (ii) the sensitivity of the T_1_w/T_2_w ratio to the latter microstructural processes, we hypothesised that the two groups would show significantly different changes in the T_1_w/T_2_w ratio. To reduce the potential for confounding factors, we monitored a number of *Zeitgeber* signals, such as food intake, caffeine intake, physical activity and exposure to blue-emitting light. To assess potential functional correlates of cortical changes, we also explored associations between the T_1_w/T_2_w ratio, and sleepiness and lapses in attention.

## 2 Methods and Materials

### 2.1 Ethics statement

This study was approved by the Regional Committee for Medical and Health Research Ethics, South-Eastern Norway (REK Sør-Øst, ref: 2017/2200) and conducted in line with the Declaration of Helsinki of 2013. All participants gave their written informed consent prior to participation and received NOK 1000.

### 2.2 Participants

The recruitment procedure was described in detail previously[18]. Volunteers were recruited through social media and a national newspaper advert. 127 volunteers underwent clinical screening over the phone and 41 were excluded due to meeting one of the following exclusion criteria: history or presence of any psychiatric disorder by means of screening questions from MINI neuropsychiatric inventory [25]; severe or chronic somatic disorder by a slightly modified version of Stanley Foundation Entry Questionnaire, which covers 21 somatic including neurological disorders and history of head injury [26]; current intake of any regular medication, smoking and caontraindications to MRI or living more than one hour of travel away from the MRI facility. Of the remaining 86 volunteers, 15 withdrew their participation, and 22 were cancelled due to logistic reasons. After study start, one participant was excluded due to illness and two due to claustrophobic reaction in the scanner. An additional five were excluded due to incomplete T_2_w data. The final sample consisted of 41 healthy adults (mean age 26 ± 6.9 years; age range 18 to 46 years; 26 women).

### 2.3 Study design

Figure 1 presents an overview of the study design. Due to time constraints and to reduce the strain on participants, T_2_-weighted MRI required for T_1_w/T_2_w ratio estimations were obtained only at the first and last MRI scan. Participants underwent the first MRI scan at time point 1 (TP1) in the morning of the first day at Oslo University Hospital, Rikshospitalet. They arrived fasting after a night of regular sleep in their own home (around 9 AM). After spending the day at the hospital, participants completed their second MRI scan (around 8 PM) and were randomised by draw to either go home to sleep (NSW group) or to stay awake at the hospital during the night (sleep deprivation group). In the morning, the NSW group returned, and both groups underwent their third MRI scan (around 8 AM). They then spent a second day at the hospital. The final scan took place in the afternoon after approximately 32 hours since study start (around 4 PM, TP2).

**Figure 1.**
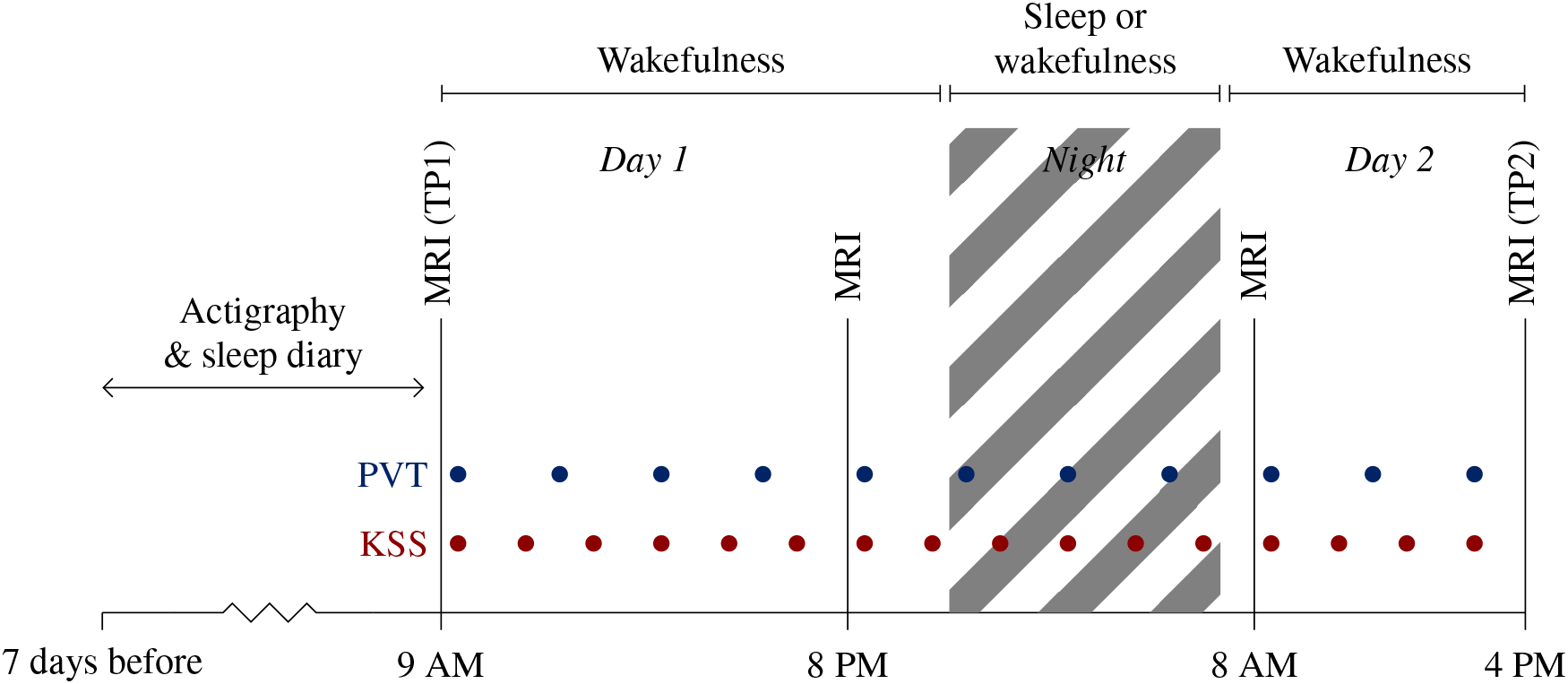
Overview of the study protocol. Participants underwent MRI including T1w- and T2w scans in the morning after a night of sleep (time point (TP)1) and then in the afternoon on the second day (TP2) after 32 hours of either normal sleep-wake or total sleep deprivation. They also underwent MRI (not including T1w- and T2w scans) in the evening on the first day and in the morning on the second day. During the study, participants underwent tests of subjective sleepiness (Karolinska Sleepiness Scale, KSS) and objective alertness (Psychomotor Vigilance Task, PVT) every second and third hour, respectively. Seven days prior to the first MRI scan, participants underwent measurements of sleep habits by actigraphy and self-report sleep diary.

A blood sample was obtained from each participant after each scan for analysis of haematocrit, to estimate hydration level. Participants followed a standardised activity plan in the company of a research assistant (see Voldsbekk et al., 2020 for a detailed description). To ensure that no one fell asleep during the MRI scans, a camera (Model 12M-i, MRC Systems GmbH, Heidelberg, Germany) inside the scanner bore was used to monitor the eyes of the participants.

### 2.4 Assessment of sleep habits

Seven days prior to study start, participants recorded their sleep pattern by self-report and actigraphy. Participants in the NSW group also recorded their sleep pattern during the night of the study. Self-report data was recorded each day by a 10-item semi-structured sleep diary[27], which assessed sleep-related behaviour and quality. The scale was modified to also include two items on caffeine intake and nicotine intake. Actigraphy data was recorded by a Condor Instruments ActTrust actigraph (São Paulo, Brazil), which measured an individual’s movements by a digital tri-axial accelerometer with a 60s epoch. In addition, participants completed five standardised questionnaires regarding their sleep habits: the Bergen Insomnia Scale[28], the Epworth Sleepiness Scale[29], the Pittsburgh Sleep Quality Index[30] and the Horne-Østberg Morningness Eveningness Questionnaire[31]. These questionnaires measure insomnia-related symptoms, daytime sleepiness, sleep quality, and chronotype, respectively.

### 2.5 Assessment of acute sleepiness and alertness

After each scan and every other hour throughout the study, participants completed the Karolinska Sleepiness Scale (KSS), which is a 1-item self-report rating of sleepiness on a nine-point Likert scale[32]. To measure objective alertness, participants completed a computerised psychomotor vigilance task (PC-PVT)[33] after each scan and every third hour of the study. For ten minutes, a five-digit millisecond (ms) counter was presented as the visual stimulus on a screen with random intervals. Participants clicked a mouse button in response. Performance was quantified as number of minor attentional lapses, i.e. reaction times (RT) > 500 ms, which has been found to be a more sensitive measure of sleep deprivation-related alertness compared to mean or median RT[34]. The task was run using Matlab 2017a (MathWorks, Massachusetts, USA) on a Lenovo laptop V510-15IKB with Windows 10 Pro and a Cooler Master mouse model SGM-1006-KSOA1. The laptop display had a refresh rate of 60 Hz.

### 2.6 MRI acquisition

Imaging was performed on a 3T Siemens Magnetom Prisma scanner (Siemens Healthcare, Erlangen, Germany) using a 32-channel head coil. The scan protocol consisted of a four-echo T_1_w multi-echo magnetisation-prepared rapid gradient-echo sequence (repetition time/echo times = 2530ms/1.69-3.55-5.41-7.27ms, field-of-view = 256×256mm^2^, voxel size = 1.0×1.0×1.0mm^3^, flip angle = 7°, acceleration factor = 2 acquisition time = 6 min, 3 seconds) and a bandwidth-matched T_2_w sequence (repetition time/echo time = 3200ms/563ms, field-of-view = 256×256mm^2^, voxel size = 1.0×1.0×1.0mm^3^, acceleration factor = 2 acquisition time = 4 min 24 seconds). The four T_1_w images were used to estimate a root mean square image which was submitted to further analyses[35].

### 2.7 MRI preprocessing

T_1_w/T_2_w maps were created using the Human Connectome Project (HCP) processing pipeline (https://github.com/Washington-University/Pipelines)[36], including processing with FreeSurfer version 5.3 (http://surfer.nmr.mgh.harvard.edu), similarly to the processing done by Grydeland and colleagues[37]. The T_1_w volume was divided by the aligned[38] and spline interpolated T_2_w volume, yielding a T_1_w/T_2_w ratio volume. For regional T_1_w/T_2_w maps, a multimodal parcellation was utilised which divided each cerebral hemisphere into 180 regions[39]. T_1_w/T_2_w values were sampled from the white matter/grey matter boundary, at three cortical depths around the middle of the cortical mantle (30%, 50%, and 70% into the cortex).

### 2.8 Statistical analyses

In order to ensure homogeneity of the sample, *t*-tests of group differences were run for each demographic and sleep-wake characteristic. In addition, we ran a *t-*test across groups to ensure that participants had a normal sleep routine at the outset of the experiment. We did not expect any group differences here, as the participants were not randomly assigned into one of the experimental groups until 12 hours into the experiment, i.e. after the second scan in the evening. To test for group differences in the T_1_w/T_2_w ratio between TP1 and TP2, the symmetrised percentage change (SPC) was calculated, which has been shown to be more robust than percentage change[40]:

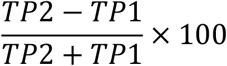

Group comparison was then run using linear regression models in R[41] in each of the 360 atlas regions. To reduce potential partial volume effects[42], the main analysis was run using the T_1_w/T_2_w ratio measured at a 50% depth. As a sensitivity analysis to assess the potential effects of cortical depth, we then reran the analysis using the T_1_w/T_2_w ratio from 30% and 70% into the cortex. As another sensitivity analysis, we re-ran our main analysis of group differences in the changes over time, using the difference between time points as outcome metric (T_1_w/T_2_w at TP 2 - T_1_w/T_2_w at TP1). The Euler number, which has been shown to be a useful index of head movement in young adults[43], was extracted from each T_1_w volume and included as a covariate. To control for multiple comparisons, we ran a custom permutation-based cluster size correction analysis across ROIs in R. A cluster was defined as adjacent regions, i.e. regions sharing a border, showing *p*-values below the cluster-forming threshold of *p* = .05. This threshold was chosen to maximise sensitivity in the smaller and less noisy parcellated brain map (compared to a voxel-wise map). To build a null model, we then ran 5000 permutations repeating the group analysis, re-shuffling the group membership for each run, and counting the maximum cluster size. Then, the size of the actual clusters was compared with this distribution of 5000 maximum cluster size from random group orderings. As a measure of effect size we calculated Hedge’s g[44] of the group difference in the means of change over time using the *effectsize* packages in R[45]. For simplicity of interpretation, as there was no difference in head movement between groups, this effect measure was calculated without head movement as a covariate (in contrast to the actual analysis).

For significant clusters, we then extracted the mean T_1_w/T_2_w ratio to perform sensitivity analyses. First, we assessed whether the group effect remained significant when including mean cortical thickness across the cluster, and haematocrit, an index of hydration. Second, to explore the functional relevance of changes in T_1_w/T_2_w ratio from TP1 to TP2, interaction analyses were performed between changes in T_1_w/T_2_w ratio and changes in sleepiness and alertness, as measured by KSS and minor lapses on the PC-PVT. To examine whether our choice of vigilance measure (minor lapses) influenced the results, we re-ran parts of the analysis using alternative measures of vigilance, namely median reaction time (RT) and RT variability (defined as the variance in RT). To correct for the multiple tests, the resulting *p*-values were adjusted by applying false discovery rate correction (FDR)[46]. We also utilised neurosynth.org, a data-driven tool for meta-analysis of the large primary literature on task-related functional MRI[47], to identify functions most implicated in the observed changes. The top 25 terms were extracted as a word cloud in which the size and colour saturation of the words correspond to the frequencies associated with each term.

## 3 Results

### 3.1 Sleep pattern assessment

As seen in Table 1, there were no differences in demographics or sleep-wake characteristics between groups. All participants had slept approximately seven hours each night for the past week, as recorded by self-report and corroborated by actigraphy measurements. For the NSW group, there was no significant difference in total sleep time on the night between the first and second day of the study compared to the night prior to study start (*t* = −.8, *p* = .42). Across groups, participants slept significantly shorter on the night before the study compared to their weekly average (*t* = 2.15, *p* = .03).

**Table 1.** Descriptive statistics for sleep-wake characteristics and tests of differences between sleep deprived and normal sleep-wake groups for each characteristic.

| | Sleep-wake<br>($n = 23$) | Sleep deprived<br>($n = 18$) | T-test of group differences |
| --- | --- | --- | --- |
| Age | 25.7 (6.97) | 26.28 (7.05) | -.26 (df = 36.46, $p = .79$, $g = -.08$) |
| Sex (no. of women) | 15 (65%) | 11 (61%) | .26 (df = 36.14, $p = .79$, $g = .08$) |
| PSQI (0 - 42)<sup>a</sup> | 4.5 (1.92) | 4.78 (1.86) | -.46 (df = 36.85, $p = .65$, $g = -.14$) |
| MEQ (16 - 86)<sup>a</sup> | 50.52 (9.16) | 50.11 (8.02) | .15 (df = 37.8, $p = .88$, $g = .05$) |
| BIS (0 - 42)<sup>a</sup> | 7.25 (5.25) | 6.39 (4.1) | .58 (df = 37.94, $p = .56$, $g = .18$) |
| ES (0 - 24)<sup>a</sup> | 6.55 (4.48) | 5.72 (2.63) | .72 (df = 34.79, $p = .47$, $g = .21$) |
| Mean TST week before study (h.hh)<sup>1,a</sup> | 7.02 (0.93) | 6.9 (0.94) | .41 (df = 36.33, $p = .68$, $g = .13$) |
| Mean TST night before study (h.hh)<sup>1,b</sup> | 6.42 (0.79) | 6.54 (1.31) | -.34 (df = 25.64, $p = .74$, $g = -.11$) |
| Mean TST night of study (h.hh) | 6.63 (0.78) | - | - |
| Change in KSS (0 - 28 h into study)<sup>c</sup> | -.57 (1.5) | 2.18 (2.96) | -3.63 (df = 23.61, $p = .001$, $g = .42$) |
| Change in PVT (0 - 30 h into study)<sup>d</sup> | 1.43 (3.34) | 3.94 (6.45) | -.9 (df = 27.88, $p = .35$, $g = .28$) |

*Note.* Mean (standard deviation). PSQI; Pittsburgh Global Sleep Quality Index. MEQ; Horne-Östberg Morningness Eveningness Questionnaire. BIS; Bergen Insomnia Scale. ES; Epworth Sleepiness Scale. TST; total sleep time. KSS; Karolinska Sleepiness Scale. PVT; psychomotor vigilance test. df: degrees of freedom. $p$ ; $p$ -value. $g$ ; Hedges $g$ .
<sup>1</sup> Reported total sleep time as measured by actigraphy and controlled against sleep diary.
<sup>a</sup> Missing for one participant.
<sup>b</sup> Missing for five participants.
<sup>c</sup> Missing for four participants.
<sup>d</sup> Missing for eleven participants.

### 3.2 Effect of 32 hours of sleep deprivation and normal sleep-wake on cortical microstructure

The change from TP1 to TP2 did not differ between groups for hydration or in-scanner head movement (*p* = .1 and *p* = .57, respectively, see Figure S1). As shown in **Figure 2A-B**, the group comparisons revealed four cortical clusters exhibiting significant differences in T_1_w/T_2_w ratio change from TP1 to TP2, spanning 37% of the ROIs in the right hemisphere (RH) and 16% in the left hemisphere (LH). Specifically, across clusters, the sleep deprived group showed an increase in T_1_w/T_2_w ratio, while the NSW group showed a decrease. In the RH, there were 66 regions across two clusters (61 and 5 regions, df=38, *p* < .0001 and *p =* .004, Hedge’s *g* = 0.83 and 0.74, respectively) in predominantly the insular, cingulate, parietal, and superior temporal cortices, while in the LH there were 28 regions across two clusters (13 and 15 regions, df=38, *p* < .0001 for both, Hedge’s *g* = 0.81 and 0.77, respectively) predominantly in the insular, cingulate, superior temporal and medial frontal cortices. These results indicate that the within-subject change between the two MRI scans were significantly different between the sleep deprivation group and the NSW group.

**Figure 2.**
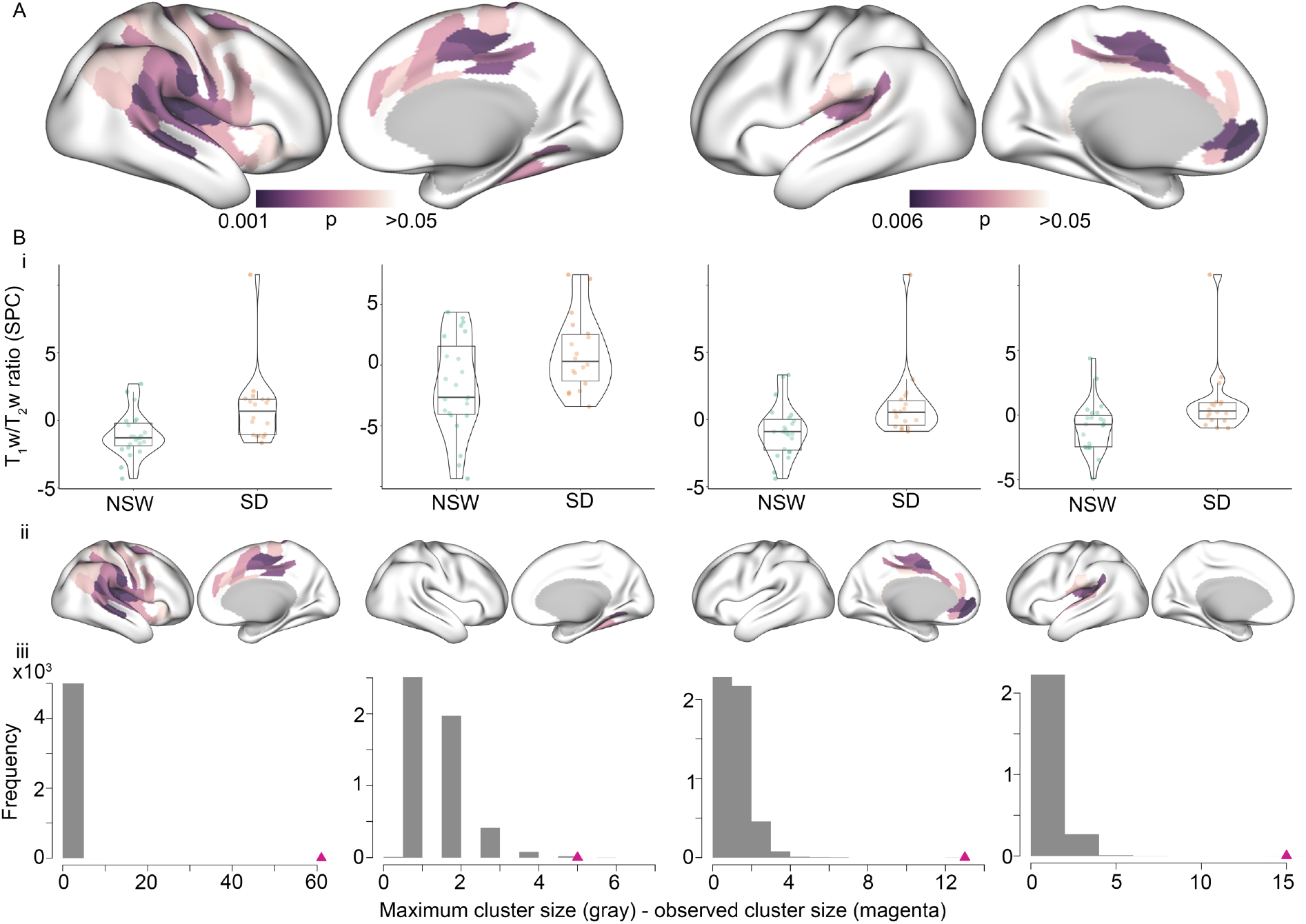
Altered intracortical T1w/T2w ratio after 32 hours of either normal sleep-wake (NSW) or sleep deprivation (SD). **A**. Surface maps showing the regions of the four clusters showing significant group differences in symmetrised percentage change (SPC) in T1w/T2w ratio in the right hemisphere between the SD group and NSW group. Each cluster was p < 0.05, and the *p* values for each region (before cluster-correction) are presented for illustration. **Bi**. Violin-box-scatter plots for the mean T1w/T2w ratio per cluster for participants in each group. **Bii**. Surface maps of each cluster. **Biii**. The distribution of maximum cluster sizes across random groupings (grey) with the actual cluster size for each cluster identified with the triangle (magenta).

The first sensitivity analysis showed that the group difference in each cortical cluster remained significant when controlling for hydration and cortical thickness (*p* < .05), except for the smallest cluster with the initially lowest effect (5-region RH cluster, *p* = .55). To assess whether the results were unduly influenced by one participant in the sleep deprivation group showing high cluster values in three of four clusters (see **Figure 2Bi**), we excluded this person was and reran the group comparison. As shown in **Figure S2**, the results were similar to the main analysis, with four clusters, comprising 77 and 14 regions in right and left hemisphere, respectively, showing group differences. The Hedge’s *g* in the RH clusters were −0.96 and −0.70, while in the LH −0.83 and −0.84, respectively.

When using a raw difference score as the outcome metric (T_1_w/T_2_w at TP 2 - T_1_w/T_2_w at TP1), two clusters showed significant differences in change over time between groups, comprising a total of 64 regions (36%) in the RH, and two clusters comprising a total of 33 regions (18%) in the LH. When excluding the one potential outlier, the corresponding results showed two clusters of 78 regions (43%) in the RH, and three clusters of 20 regions (11%) in the LH. As shown in **Figure 3,** there was regional variation in the strength of the effect specific to each group. Specifically, the NSW group showed a stronger reduction across medial parietal regions, fusiform gyrus, and the posterior cingulate cortex, while the sleep deprived group showed a stronger increase in the anterior cingulate cortex, as well as both lateral and medial frontal regions. As determined by neurosynth.org[47] (**Figure 4**), the observed changes implicated regions most strongly involved in attention, listening, movement and pain.

**Figure 3.**
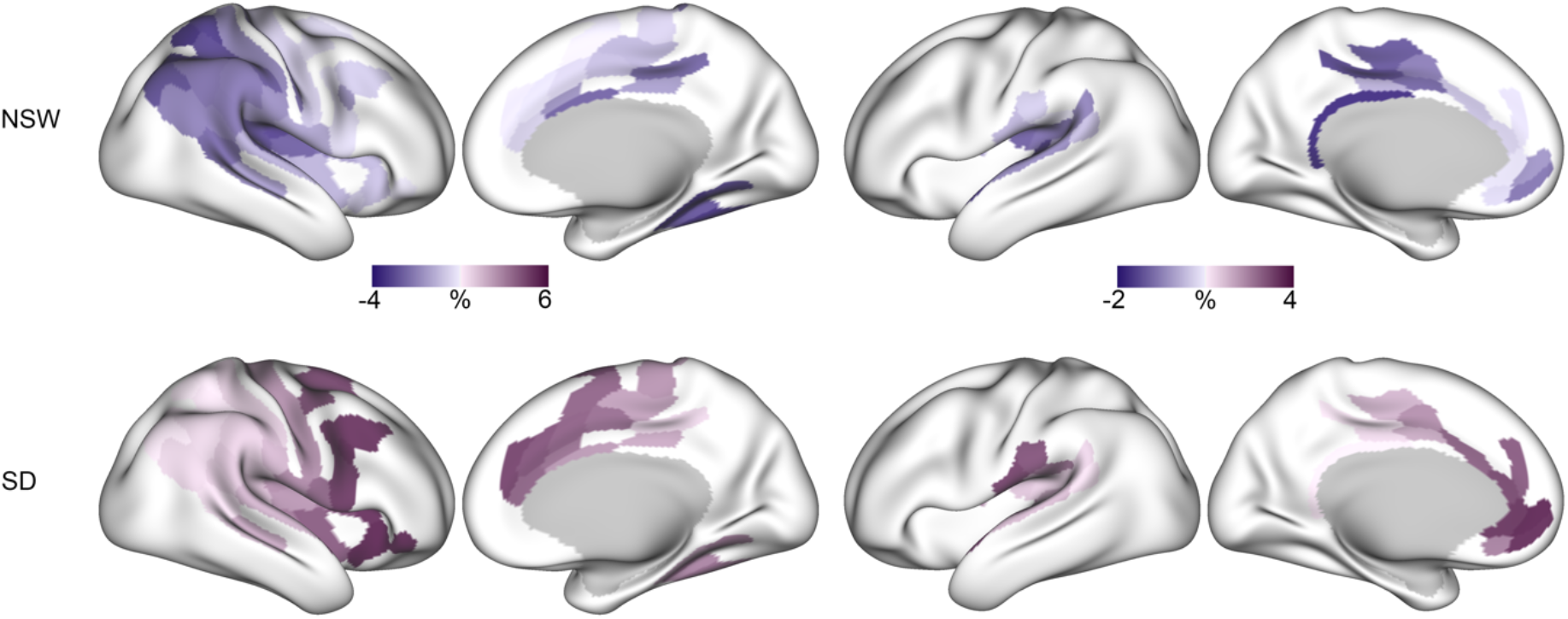
Regional variation in sensitivity to either normal sleep-wake (NSW) or sleep deprivation (SD). Percentage change within each significant cluster for the NSW group (top) and SD group (bottom).

**Figure 4.**
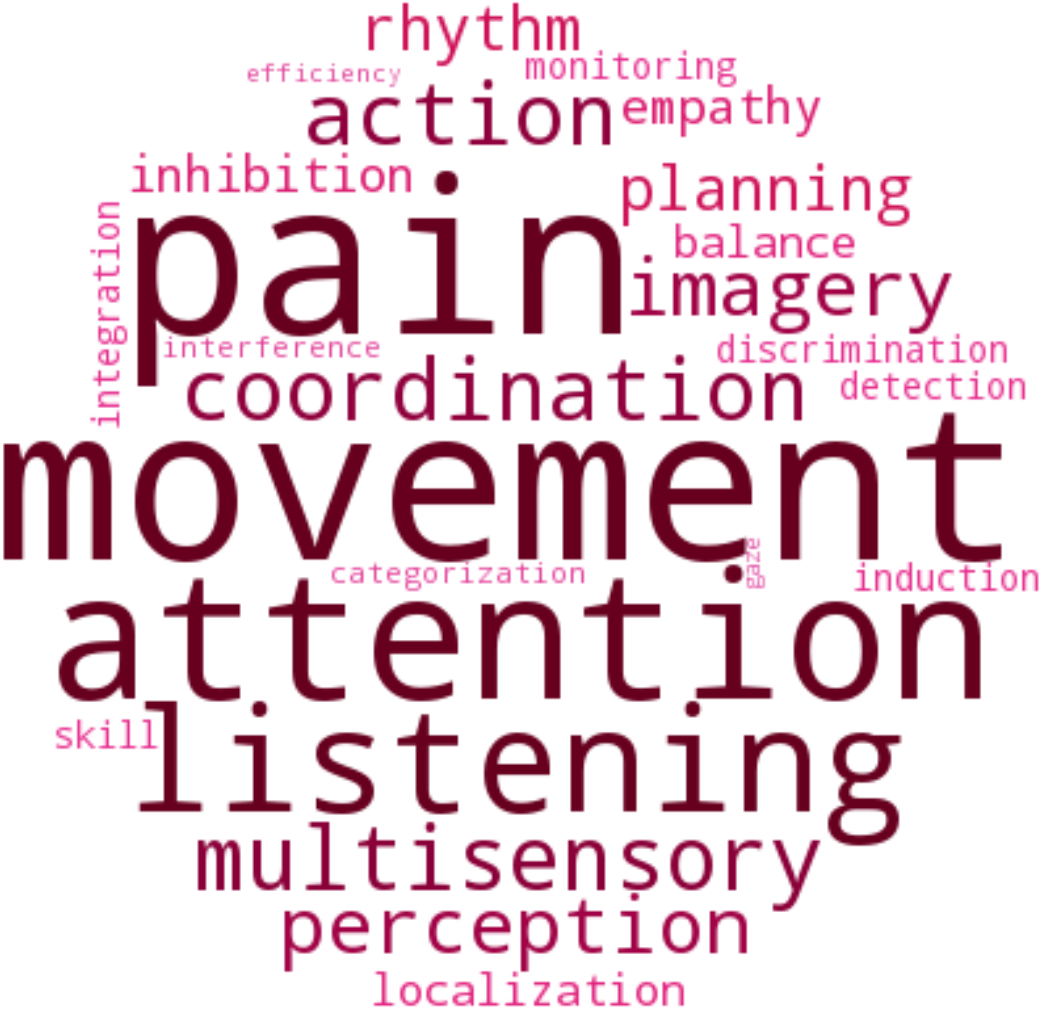
Word cloud based on correlations with NeuroSynth meta-analysis maps across groups. Larger size and colour saturation indicate stronger correlation.

To assess whether the higher number of RH regions showing group differences was partly due to a thresholding effect, we summarised the *p*-values of the regions in the LH which were only significant in the RH. Of these 38 LH regions, the median *p*-value was .10 (minimum – maximum is .02 - .55), indicating that a portion of the LH regions likely would have been significant with higher statistical power.

The sensitivity analyses assessing the effects of cortical depth, shown in **Figure 5**, revealed significant group differences using T_1_w/T_2_w ratio values extracted from 30% and 70% cortical depth. At 30% depth, a slightly higher number of regions significantly differed between groups (97 RH regions and 29 LH regions), as compared with the results from 50% depth, while a slightly lower number of significant regions were found at 70% depth (47 RH regions and 14 LH regions).

**Figure 5.**
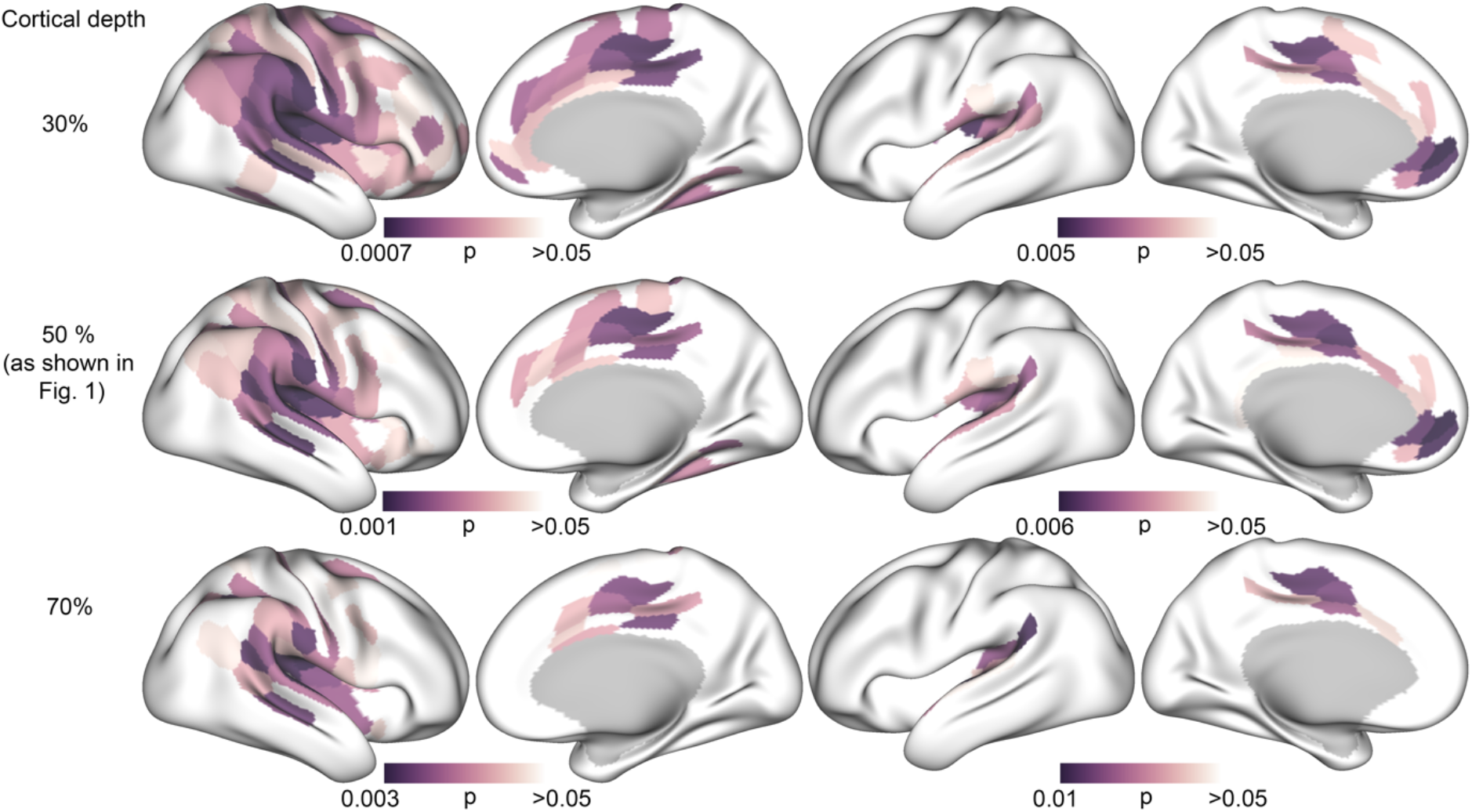
Surface maps showing significant group differences in T1w/T2w ratio in the right hemisphere (similar to results reported in Figure 2A) as a function of cortical depth. Top row: T1w/T2w ratio sampled from 30% into the cortex from the white/grey matter boundary. Middle row: 50% cortical depth (as shown in Figure 2A). Bottom row: 70% cortical depth.

### 3.3 Associations between changes in cortical microstructure and sleepiness and attention

The difference in subjective sleepiness and in objectively measured attentional lapses over the 32 hours are presented in Figure S3, and the distribution of change values in Figure S4. The change in subjective sleepiness differed between the groups (see Table 1), with the sleep deprived group showing increased sleepiness (higher KSS at TP2). There was no group difference in change in lapses in attention. Alternative measures of attention from the PVT, namely median RT and the variance of RT, showed similar group differences in alertness (see Figure S5).

The interaction analyses of an association between T_1_w/T_2_w ratio changes and sleepiness and attentional lapses, did not show any significant results after FDR-correction (*p* > .28, these analyses were run on the results excluding the participant with a large change value). The uncorrected results showed a relation between changes in T_1_w/T_2_w ratio and sleepiness (see Figure S6), with an effect in the 9-region LH cluster (*p* = .04), while the other clusters did not show effects (*p* = .61 for RH cluster 1, *p* = .56 for RH cluster 2, and *p* = .25 for LH cluster 2). The effect was due to a negative correlation between T_1_w/T_2_w change and sleepiness change in the sleep deprived group (*r* = −.7, *p* = .003), while the NSW group showed an opposite, and weaker relation (*r* = .27, *p* = .27). Thus, in the sleep deprivation group, a stronger increase in T_1_w/T_2_w ratio was related to lower subjective sleepiness.

The uncorrected results of changes in T_1_w/T_2_w ratio and lapses in attention (see Figure S7) showed an interaction in the 4-region RH cluster (*p* = .022), while the other clusters did not show effects (*p* = .43 for RH cluster 1, *p* = .43 for LH cluster 1, and *p* = .36 for LH cluster 2). This trend was due to a positive correlation between T_1_w/T_2_w change and alertness change in the NSW group (*r* = .67, *p* = .012), while the sleep deprivation group showed a weaker relation (*r* = −.16, *p* = .54). The results using alternative measures of vigilance were similar, with only the the 4-region RH cluster showing significant (variance RT *p* = .01) or near-significant results (median RT *p* = .059).

## 4 Discussion

The findings of the present study indicate that 32 hours of sleep deprivation yields different intracortical changes in the T_1_w/T_2_w ratio than compared to an NSW cycle. The differences between the sleep deprivation and the NSW groups were more prominent in the RH, but were observed in both hemispheres in insular, cingulate and superior temporal cortices and included regions involved in attentional, auditory, movement and pain processing[47]. Across regions, the sleep deprived group showed an increased T_1_w/T_2_w ratio, whereas the NSW group exhibited a reduced ratio. The effects were not explained by an estimate of in-scanner head movement, were present at various depths in the cortex, and 95% of the effects remained significant after adjusting for cortical thickness and hydration. Although speculative, candidate neurobiological mechanisms of the observed cortical effects include myelin and dendrite density, previously linked to the T_1_w/T_2_w ratio[14-16].

We found group differences in T_1_w/T_2_w ratio change in the bilateral cingulate, insula, and superior temporal cortices and in right parietal and left middle frontal regions. Although this is the first study of T_1_w/T_2_w ratio in sleep deprivation, these results overlap with findings from three previous studies. The first study reported reduced cortical volume mainly in the insular, parietal, posterior cingulate, motor, and somatosensory cortices after 36 hours of sleep deprivation[8]. The second study found increased grey matter density in the frontal pole and the superior and middle frontal gyri, as well as decreased volume and thickness in the temporal pole after 24 hours of sleep deprivation[48]. The third study reported reduced thickness in bilateral medial parietal regions, yet did not detect a significant group by time interaction effect when compared to a NSW group[9]. Thus, the results of the present and previous studies suggest that acute sleep deprivation is associated with cortical alterations, mainly in frontal, temporal, parietal, and insular cortices. The current study points to new effects in the bilateral anterior cingulate, the superior temporal (including auditory cortex), and the left medial frontal regions. This difference in results may reflect the differences in study designs, but also indicate that the microstructural T_1_w/T_2_w ratio has greater sensitivity to the intracortical effects of wake and sleep deprivation.

We observed increased T_1_w/T_2_w ratio in the sleep deprived group. Further histological studies are needed to elucidate the neurobiology underlying these T_1_w/T_2_w ratio changes, yet they may be linked to wake-related processes, circadian rhythm mechanisms, or a combination of the two[49]. Although the precise neural effects of wake and sleep remain to be clarified, the synaptic homeostasis hypothesis posits that wake and sleep are associated with net increases and decreases, respectively, in densities of synapses and dendrites[50]. Consistent with this hypothesis, spontaneous and enforced wake in animals were linked to increased dendritic branching and synapse number[6, 51, 52] and reduced synapse pruning[53]. Thus, although speculative, T_1_w/T_2_w ratio changes within the sleep deprivation group of the present study could be related to wake-induced increases in synaptic and dendritic densities.

An alternative explanation for the increased T_1_w/T_2_w ratio after sleep deprivation is cortical myelin changes. The T_1_w/T_2_w ratio correlates with cortical myelin content[14], and the ratio was higher in myelinated than demyelinated cortex in multiple sclerosis[15]. There is a scarcity of studies of cortical myelin after sleep deprivation in humans, yet two previous MRI studies reported RD reductions in human white matter after extended wake[10, 18]. Reduced RD is, in general, associated with increased myelination[12]. Furthermore, a growing body of animal data links sleep and wake to myelin and oligodendrocyte alterations[7, 54]. Thus, further studies should clarify whether changes to myelin and oligodendrocytes contribute to the T_1_w/T_2_w ratio increases found in the current study.

We observed a T_1_w/T_2_w ratio reduction within the NSW group from the morning of the first study day to the afternoon of the second day. Several lines of evidence point to sleep-wake as an active process throughout both night and day, and potentially involving alterations in brain microstructure. Two separable, yet interacting processes are considered to regulate sleep-wake: sleep homeostasis, by which increasing sleep pressure accumulates as a function of time spent awake, and the circadian rhythm, which is an intrinsic oscillating cycle of 24 hours regulated by exposure to daylight[49]. In line with this framework, diurnal fluctuation has been observed in the transcription of genes related to macromolecule homeostasis[55, 56], oligodendrocyte proliferation, phospholipid synthesis and myelination[7]. As such, we speculate that T_1_w/T_2_w changes from morning to evening of the first study day would have been reversed by the night of sleep, and that the NSW group changes detected here reflect morning- to-afternoon processes of the second study day. Importantly, these hypotheses must be tested in future studies since the current work obtained T_2_-weighted MRI only at baseline (TP1) and after 32 hours (TP2). Notwithstanding this limitation, the T_1_w/T_2_w ratio reduction from the morning of the first study day to the afternoon of the second day within the NSW group is consistent with recent studies reporting time-of-day effects on other MRI modalities[9, 10, 13, 18–24].

Taken together, the directionality of the T_1_w/T_2_w ratio changes in the two groups were, interestingly, in the opposite direction. Simultaneous MRI and histological analyses are required to clarify whether regular wake length, e.g., from morning to the same afternoon, could be associated with neural processes that qualitatively differ from those induced by sleep deprivation. The relationships between wake length and cortical alterations are not necessarily linear. In support of this notion, one study observed decreases in cortical thickness from morning to afternoon (2 hours)[21], whereas another study detected increased cortical thickness from morning to evening (14 hours)[9]. Furthermore, the T_1_w/T_2_w ratio changes observed in the current study could also reflect the likely complex and dynamic relationship between wake-induced neural processes and the circadian rhythm of the cortex. The current understanding of sleep entails two separable, yet interacting processes, namely the homeostatic process, in which sleep pressure accumulates as a function of time spent awake, and the circadian process, which oscillates at about 24-hour periods and is regulated by exposure to daylight[49].

The T_1_w/T_2_w ratio group differences comprised regions involved in attentional processes, which are amongst the first to deteriorate in response to sleep deprivation[57]. However, we found no significant relationship between attentional lapses and T_1_w/T_2_w ratio. Previous studies reported significant associations between sleepiness and changes in cortical thickness[9] and cortical density[48] after sleep deprivation. Consistent with these results, we found a weak association between reported sleepiness and T_1_w/T_2_w ratio change in one of the smaller clusters, but this did not survive FDR-correction. Thus, while the group differences suggest significant effects of sleep deprivation, the functional consequences of the intracortical T_1_w/T_2_w ratio changes among the sleep deprived individuals remain to be elucidated.

Several study limitations should be noted. First, participants had slept less the night before the study than their weekly average. However, if we assume that sleep reverses microstructural changes after wakefulness, we expect less sleep to result in attenuation, rather than inflation, of any sleep deprivation-related changes. Second, participants in the NSW group went home to sleep, which was measured with an actigraph and a sleep diary. Sleep in a laboratory with polysomnography would have resulted in greater control of exposure to *Zeitgebers* and sleep length. Third, the study did not include T_2_w MRI acquisitions from the evening of the first study day and the next morning, which limited our ability to interpret the temporal dynamics of the T_1_w/T_2_w ratio changes. Finally, the sample size was modest, and the analyses indicated that more LH effects would have been significant with higher statistical power. The lateralisation of T_1_w/T_2_w ratio effects with more significant clusters in the RH should therefore be cautiously considered. In addition, the between-subject design employed introduces additional degrees of uncertainty related to possible inter-subject differences when it comes to vulnerability to sleep deprivation. Given the relatively modest sample size, this represents an important consideration in the interpretation of our results.

The current study provides evidence that compared with a NSW cycle, 32 hours of sleep deprivation yields intracortical microstructural changes as indexed by the T_1_w/T_2_w ratio. These effects were observed within regions linked to attentional, auditory, movement and pain processing and were not explained by an estimate of in-scanner head movement, minimally affected by cortical thickness, and hydration, and were present at various depths in the cortex. The T_1_w/T_2_w ratio changes could reflect alterations in myelin or dendrite density, or both, and histological analyses are needed to clarify the precise underlying cortical processes.

## Acknowledgements

This project was funded by research grants from the Norwegian South-East Health Authorities (2018077, 2017090, 2015078), the Research Council of Norway (249795), the Centre for Digital Life Norway, the Ebbe Frøland foundation, the Norwegian Competence Center for Sleep Disorders, Haukeland University Hospital, Bergen (www.sovno.no), the University of Oslo Life Science summer scholarship for students and a research grant from Mrs. Throne-Holst.

