## supplementary material for "Evidence for widespread alterations in cortical microstructure after 32 hours of sleep deprivation"

### 1 Change in Euler number and hydration in each group

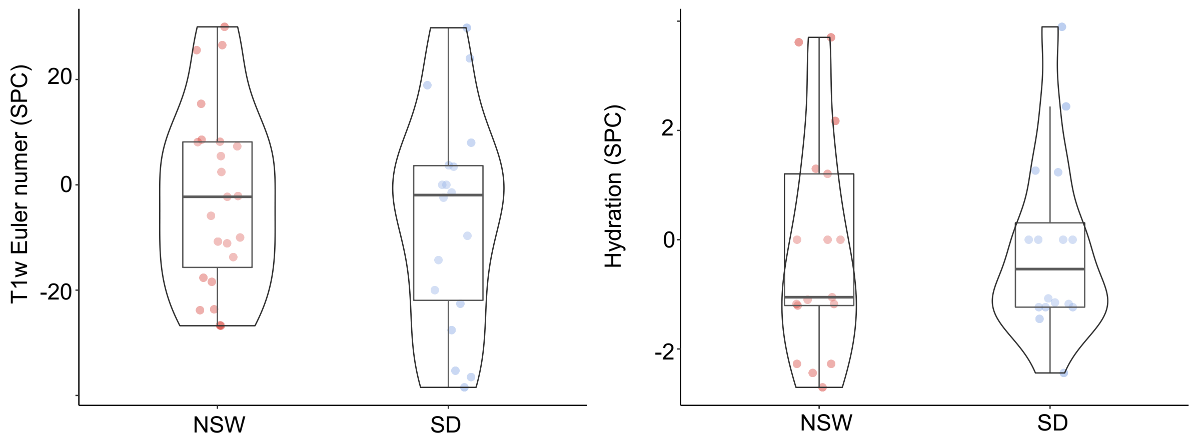


### 2 Main analysis rerun excluding one participant in the sleep deprivation group

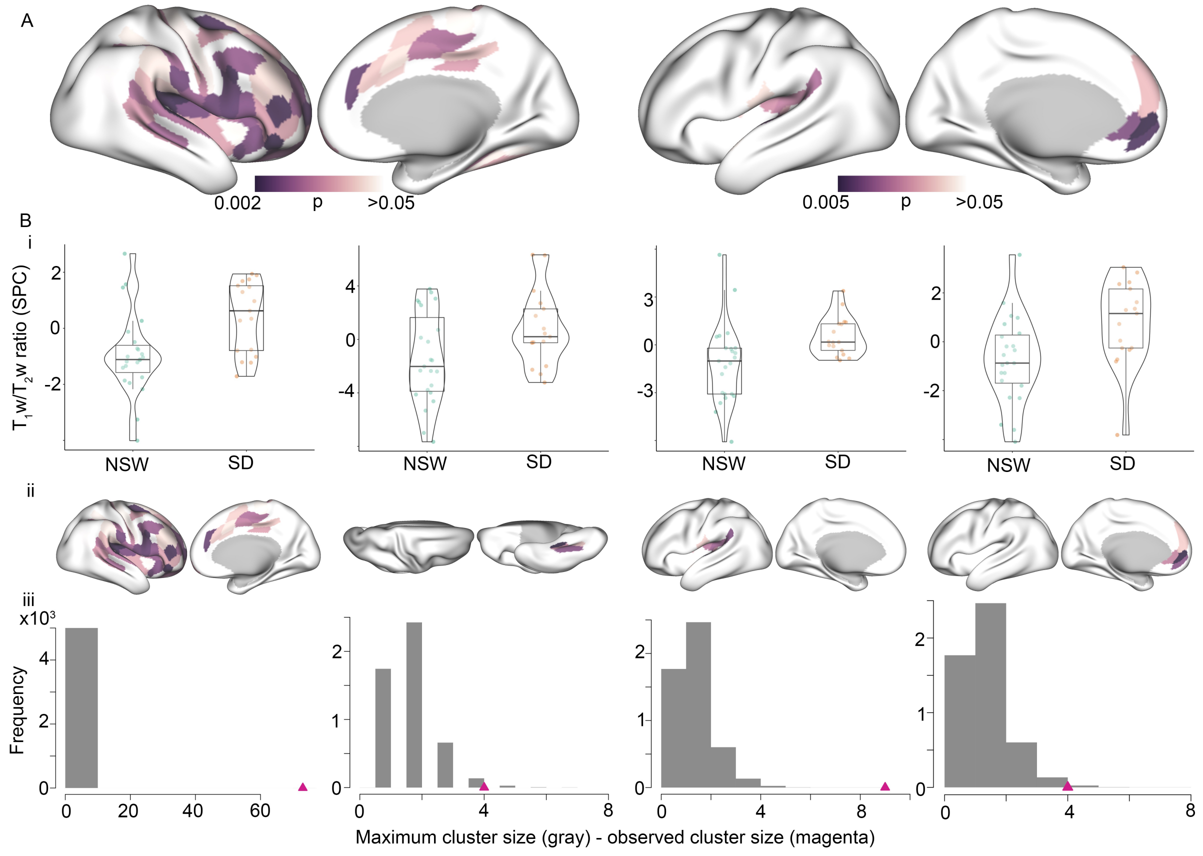

### 3 Difference in sleepiness and attention across the 32 hours of the study

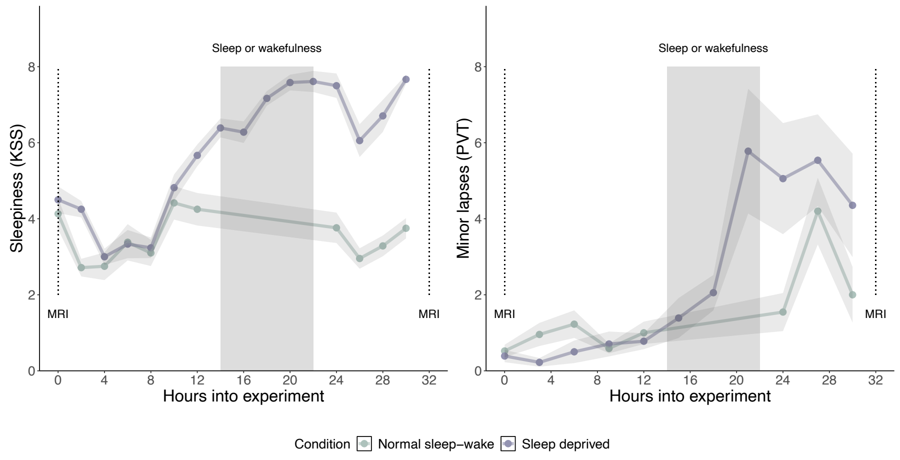


### 4 Distribution of change in sleepiness and attention from TP1 to TP2

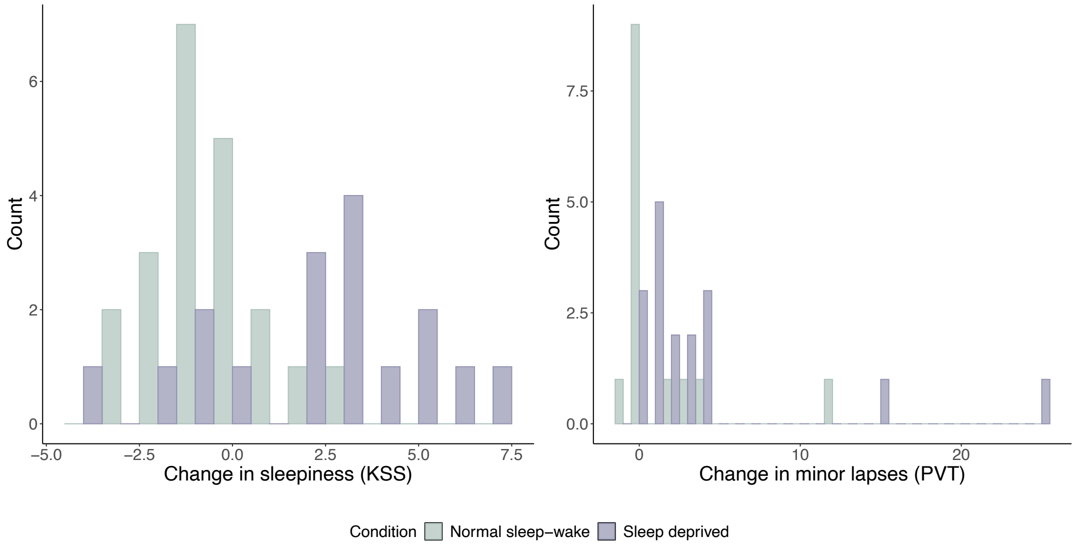


### 5 Alternative measures of attention from the PVT: group differences

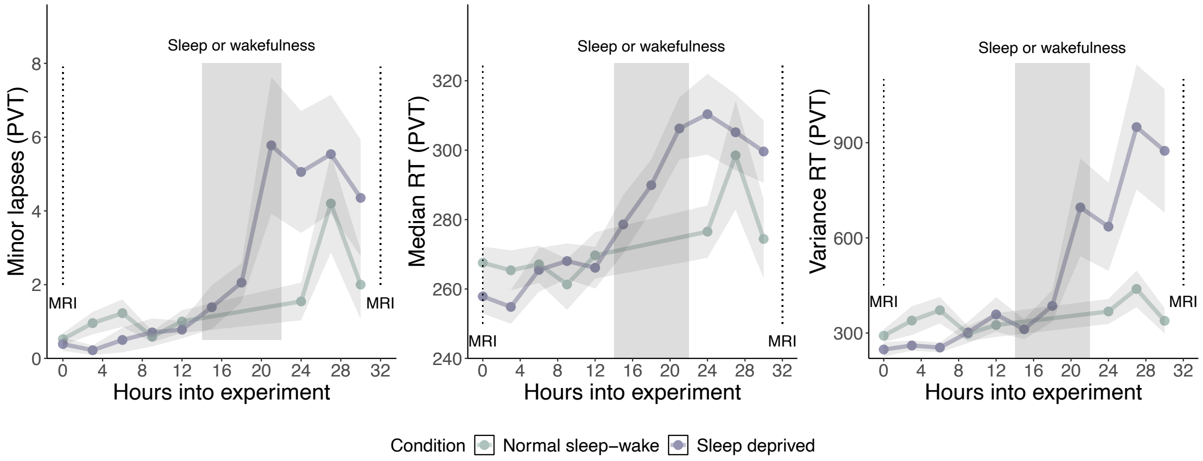

### 6 Uncorrected correlations between changes in $T_1w/T_2w$ ratio and sleepiness

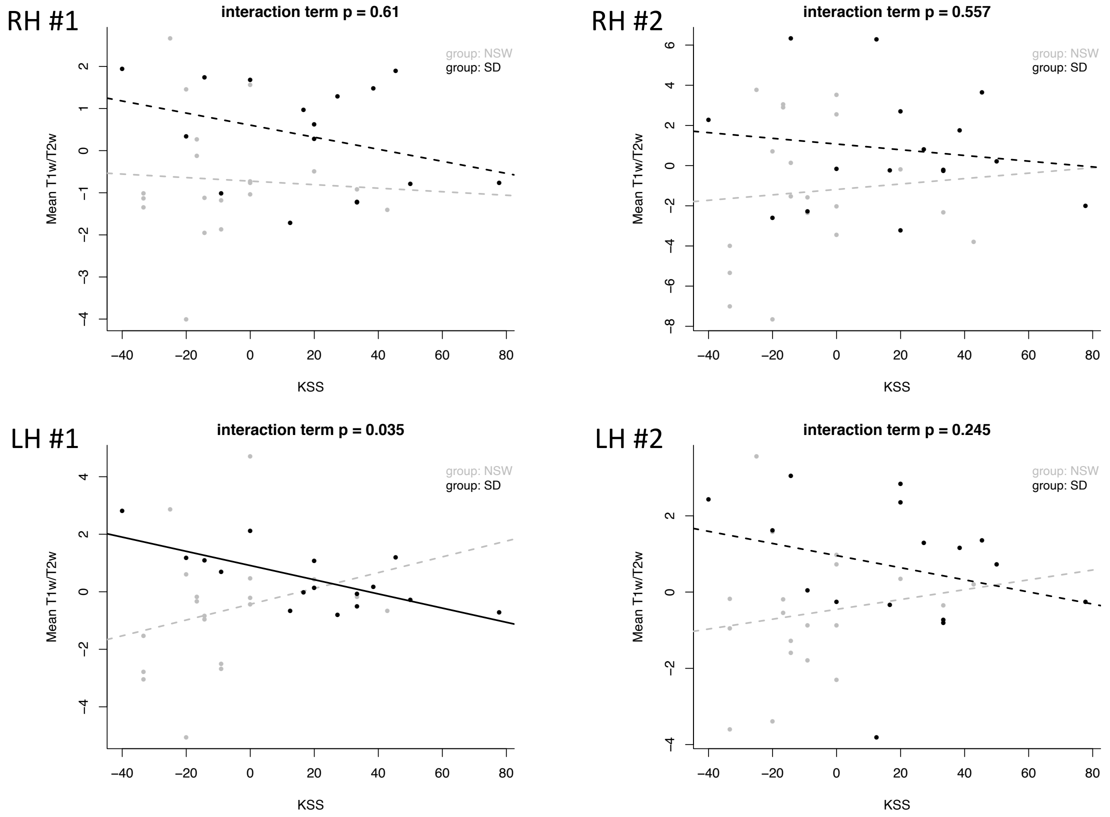


### 7 Uncorrected correlations between changes in $T_1w/T_2w$ ratio and lapses in attention

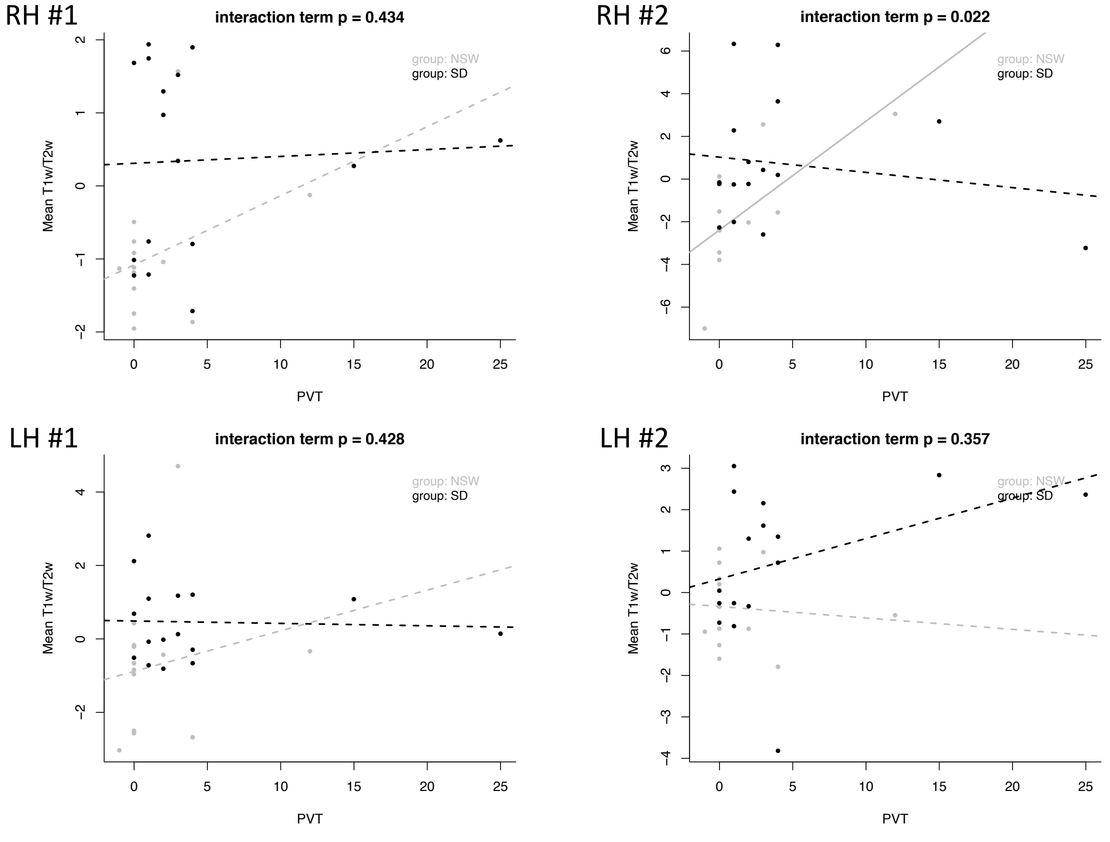
